# Vitamin D Receptor Upregulates Tight Junction Protein Claudin-5 against Tumorigenesis

**DOI:** 10.1101/2021.04.29.441977

**Authors:** Yongguo Zhang, Shari Garrett, Robert E. Carroll, Yinglin Xia, Jun Sun

## Abstract

**Background/Objective:** Tight junctions (TJs) are essential for barrier integrity, inflammation, and cancer. The TJ protein Claudin-5 in the epithelia forms paracellular barriers and pores for permeability. Vitamin D and the vitamin D receptor (VDR) play important roles in various cancers. Although VDR and Claudin-5 are all involved in colorectal cancer (CRC), it remains unclear if they are closely related or function independently.

**Design:** Using the human CRC database, we explored the correlation between VDR and Claudin-5. We then investigated the VDR regulation of Claudin-5 using VDR knockout (VDR^-/-^) and intestinal epithelial VDR knockout mice (VDR^ΔIEC^) with chemical-induced colon cancer and an epithelial VDR overexpression model. Human samples, organoids, and intestinal epithelial cells were used to determine the underlying mechanisms.

**Results:** In human colon cancer, colonic VDR expression was low and was significantly correlated with a reduction of Claudin-5 mRNA and protein. In the colon of VDR^-/-^ and VDR^ΔIEC^ mice, deletion of VDR led to lower protein and mRNA levels of Claudin-5. Intestine permeability was increased in the AOM-DSS-induced VDR^-/-^ colon cancer model. Lack of VDR and a reduction of Claudin-5 are associated with an increased number of tumors in the VDR^-/-^ and VDR^ΔIEC^ mice. Furthermore, gain and loss of function studies have identified *CLDN-5* as a downstream target of the VDR signaling pathway. Epithelial VDR overexpression protected against the loss of Claudin 5 in response to intestinal inflammation

**Conclusion:** This study advances the understanding of how VDR regulates intestinal barrier functions in tumorigenesis as a biomarker and potential treatment.

**A short summary:**

1. *What is already known about this subject?*

- Tight junction structures are essential for intestinal barrier integrity, inflammation, and cancer.
- Vitamin D deficiency and the vitamin D receptor (VDR) play important roles in the development of colon cancer.
2. *What are the new findings?*

- Our study is the first to link barrier function, a specific tight junction protein, and genetic susceptibility through intestinal epithelial VDR in human colorectal cancer.
- Our study fills an existing gap by characterizing the mechanism of intestinal epithelial VDR in regulating barrier functions through alterations in TJs in tumorigenesis. VDR is important for the maintenance of the physiological level of the TJ protein Claudin-5 in the colon. The *CLDN-5* gene is a downstream target of the VDR signaling pathway. Lack of VDR led to a reduction of Claudin-5 in tumors, whereas enhancing VDR increased Claudin-5 to protect the intestinal epithelial cells from tumorigenesis.
- We report fecal VDR reduction in a colon cancer model. This introduces the possibility for the identification of new biomarkers and therapeutic targets to restore VDR-dependent functions in CRC.
3. *How might it impact on clinical practice in the foreseeable future*

- Diagnosis of CRC considering VDR status
- Barrier: direct, indirect biomarkers
- Intestinal barriers in cancer prevention and treatment

Barrier function and VDR are not only essential for the maintenance of intestinal homeostasis, but they are also critical for the development of chronic mucosal inflammation and cancer. This knowledge can be immediately used to develop intestinal VDR and Claudin-5 as clinical biomarkers for identifying patients who may benefit from currently available interventions and could also be used for the eventual development of novel strategies for the prevention and treatment of human CRC.

## Introduction

Tight junction structures are essential for intestinal innate immunity and barrier function. The disruption of TJs is a common manifestation of various diseases, including chronic inflammation and cancer. Changes in the expression and distribution of TJ proteins such as Claudin-2, −5, and −8 lead to discontinuous TJs and barrier dysfunction in active Crohn’s disease (CD), a type of inflammatory bowel disease [1]. Claudin-5 is expressed in the epithelia and endothelia and forms paracellular barriers and pores that determine permeability. This protein is downregulated in colon cancer [2, 3].

VDR is a nuclear receptor that mediates most known functions of the biologically active form of vitamin D [4, 5, 6]. VDR possesses multiple critical functions in regulating innate and adaptive immunity, intestinal homeostasis, host response to microbiota, and tight junction structure [7, 8, 9, 10, 11, 12, 13, 14]. Vitamin D/VDR deficiency has been implicated in patients with inflammatory bowel disease and colon cancer [15, 16, 17, 18, 19, 20, 21]. Our study demonstrated that VDR is essential for maintaining intestinal and microbial homeostasis [22] and for protecting against intestinal tumorigenesis [23] [24]. Although vitamin D has been extensively studied, many critical questions regarding the biological functions of intestinal VDR in CRC remain unanswered. Although VDR and TJ proteins (e.g., Claudins) are involved in colon cancer, it remains unclear if they are closely related or function independently. Considering the multiple functional roles of VDR in the development of colon cancer [20, 24], it is important to dissect the cellular and molecular mechanisms by which VDR contributes to barrier function in protecting the host from tumorigenesis.

Here, we revisited the human CRC database and determined that colonic VDR expression is low and positively correlated with the reduction of the TJ protein Claudin-5 in CRC, including colitis-associated colon cancer. We investigated the novel role of VDR in regulating Claudin-5 expression using VDR^-/-^ and intestinal epithelial VDR knockout mice (VDR^ΔIEC^) in a colitis-associated colon cancer model. Human organoids, human colon cancer samples, VDR^-/-^ mouse embryonic fibroblasts (MEF) cells, and cultured intestinal epithelial cells were used to determine the molecular mechanisms. We determined that VDR is an important transcriptional regulator for the maintenance of physiological levels of the target gene *Claudin-5* in the intestine. Furthermore, we generated a conditional intestinal epithelial VDR-overexpressed mouse model to study the protective role of VDR in the maintenance of TJs in the context of inflammation. Our goal was to provide a detailed understanding of how VDR status contributes to intestinal inflammation and cancer. Our findings may offer an additional avenue to treat colon cancer by restoring barrier functions and developing a new protocol for risk assessment and prevention of cancer.

## Results

### Reduced VDR was positively correlated with low Claudin-5 expression in CRC patients

We first examined the gene expression levels of VDR and Claudin-5 in normal and human CRC samples by reviewing the GEO databases GSE 4183 and GSE 8671 from Affymetrix data (human genome U133 Plus 2.0 arrays). Reduced VDR and Claudin-5 expression was observed in patients with CRC (**Fig. 1A**). To quantify and visualize the correlations between intestinal Claudin-5 and the VDR protein, we performed a regression analysis of VDR against Claudin-5 and conducted a scatter plot with a regression line (**Fig. 1B**). We found significantly coordinated expression of VDR and Claudin-5 in biopsy samples collected from patients with CRC. We further analyzed data obtained from human colitis-associated colon cancer (**Fig. 1C)**. VDR and Claudin-5 expression was significantly reduced in patients with colitis-associated CRC (GEO database GSE8671, GSE10714, and GSE37283) (**Fig. 1C)**. We identified a positive correlation between VDR and Claudin-5 in biopsy samples collected from colitis-associated CRC patients and healthy controls (**Fig. 1D)**. We then examined the protein levels of intestinal VDR in normal and CRC human colon samples using IHC. Compared to normal intestines, CRC patients with CRC possessed significantly lower VDR expression (**Fig. 1E)**. Immunofluorescence (IF) staining of Claudin-5 revealed significantly lower Claudin-5 expression in CRC human colon samples (**Fig. 1F)**. We performed correlation analysis and scatter plots of the staining intensity changes between the VDR protein and Claudin-5 in the colon. The results revealed that the staining intensity of Claudin-5 and intestinal VDR was positively associated with the Pearson correlation coefficient **(Fig. 1G)**. Thus, we revealed that colonic VDR expression is low and is correlated with the reduction of Claudin-5 at both mRNA and protein levels in human CRC, including colitis-associated colon cancer.

**Fig. 1.**
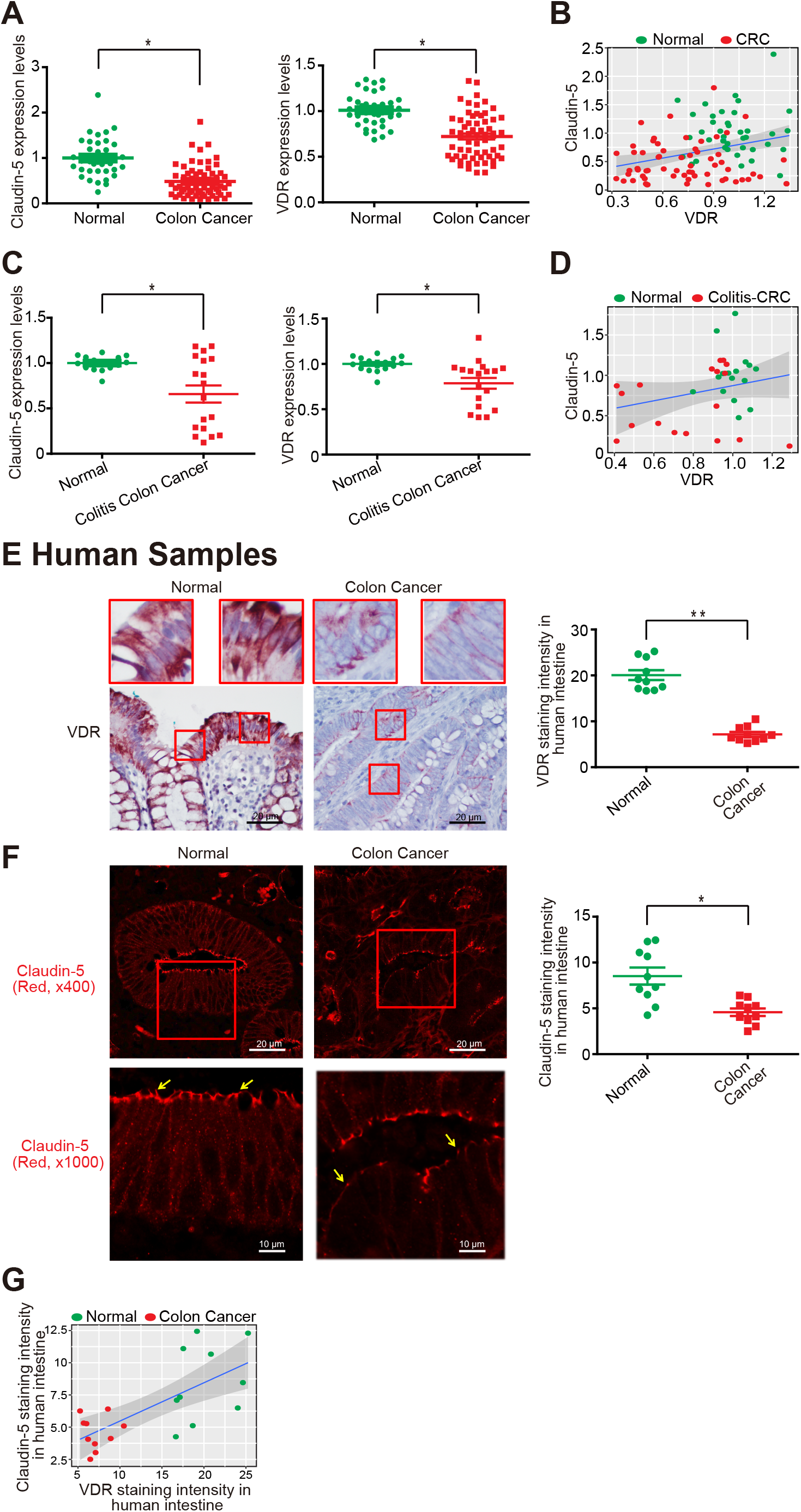
Reduced VDR was correlated with low Claudin-5 expression in human CRC patients. **(A)** Reduced VDR and Claudin-5 expression in patients with CRC (GEO database GSE4183 and GSE8671 (data were expressed as mean ± SD; Normal, n=40; CRC, n=62; student t test, * P < 0.05). **(B)** Significantly coordinated expression of VDR and Claudin-5 in biopsy samples collected from CRC patients. We performed a regression of VDR against Claudin-5 and conducted a scatter plot analysis with a regression line (GEO database GSE4183 and GSE8671, Normal, n=40; CRC, n=62; Intercept = 0.244; Slope = 0.5297). Values for healthy controls are presented in blue and values for CRC patients are presented in red. **(C)** Reduced VDR and Claudin-5 expression in patients with Colitis-associated CRC (GEO database GSE8671, GSE10714 and GSE37283 (data were expressed as mean ± SD; Normal, n=16; Colitis-associated CRC, n=18; student t test, * P < 0.05). **(D)** Coordinated expression of VDR and Claudin-5 in biopsy samples collected from Colitis-associated CRC patients. We performed a regression of VDR against Claudin-5 and conducted a scatter plot analysis with a regression line (GEO database GSE8671, GSE10714 and GSE37283 Normal, n=16; Colitis-associated CRC, n=18; the correlation is 0.2549 with p-value = 0.1457). **(E)** Intestinal VDR staining in normal and CRC human colon samples. Compared to normal intestines, intestines from CRC patients possessed significantly lower VDR expression. (Images are representative of experiments performed in triplicate; Normal, n=10; Colorectal cancer, n=10; Student *t* test; \**P* < 0.05). **(F)** IF staining of Claudin-5 in normal and CRC human colon samples. Compared to normal intestines, the intestines of CRC patients exhibited significantly lower Claudin-5 expression. (Images are representative of experiments in triplicate; Normal, n=10; Colon cancer, n=10; Student *t* test; \**P* < 0.05). **(G)** The correlation analysis of staining intensity between intestinal Claudin-5 and VDR in human colon samples. (P < 0.0734, n = 6 for Normal and Colon cancer, respectively).

### Larger and more tumors developed in VDR deficient mice

Animal models have been developed to reflect the initiation and progression of human colon cancer [25]. Azoxymethane (AOM) [26] mice develop hyperproliferative colonic mucosa, aberrant crypt foci (ACF), and eventually carcinomas [27]. An AOM–dextran sulfate sodium (DSS) model is widely used to study colitis-associated colon cancer [28]. We next investigated the role of VDR in regulating Claudin-5 expression in the development of cancer using an AOM/DSS-treated mouse model (**Fig. 2A**). For wild-type VDR^+/+^ and whole-body VDR knockout (VDR^-/-^) mice, representative colons with tumors are shown (**Fig. 2B).** We observed that AOM/DSS-treated VDR^-/-^ mice developed more tumors in the colon (**Fig. 2C**). The maximum tumor size was significantly larger in VDR^-/-^ mice compared to that in VDR^+/+^ mice (**Fig. 2D**). Furthermore, pathological analysis of colon samples indicated differences in tumor stage (carcinoma versus adenoma) between VDR^-/-^ mice and the VDR^+/+^ AOM/DSS experimental groups (**Fig. 2E**). Epithelial hyperproliferation plays a critical role in the development of cancer. The IF data of the proliferative marker PCNA revealed that PCNA in the colon was significantly increased in the VDR^-/-^ mice compared to that in the VDR^+/+^ mice (**Fig. 2F**). Chronic inflammation is one of the factors that contribute to CRC. We determined that the serum cytokines TNF-α, IL-1β, and IL-17 were significantly higher in the VDR^-/-^ mice, compared to levels in the VDR^+/+^ mice (**Fig. 2G**).

**Fig. 2.**
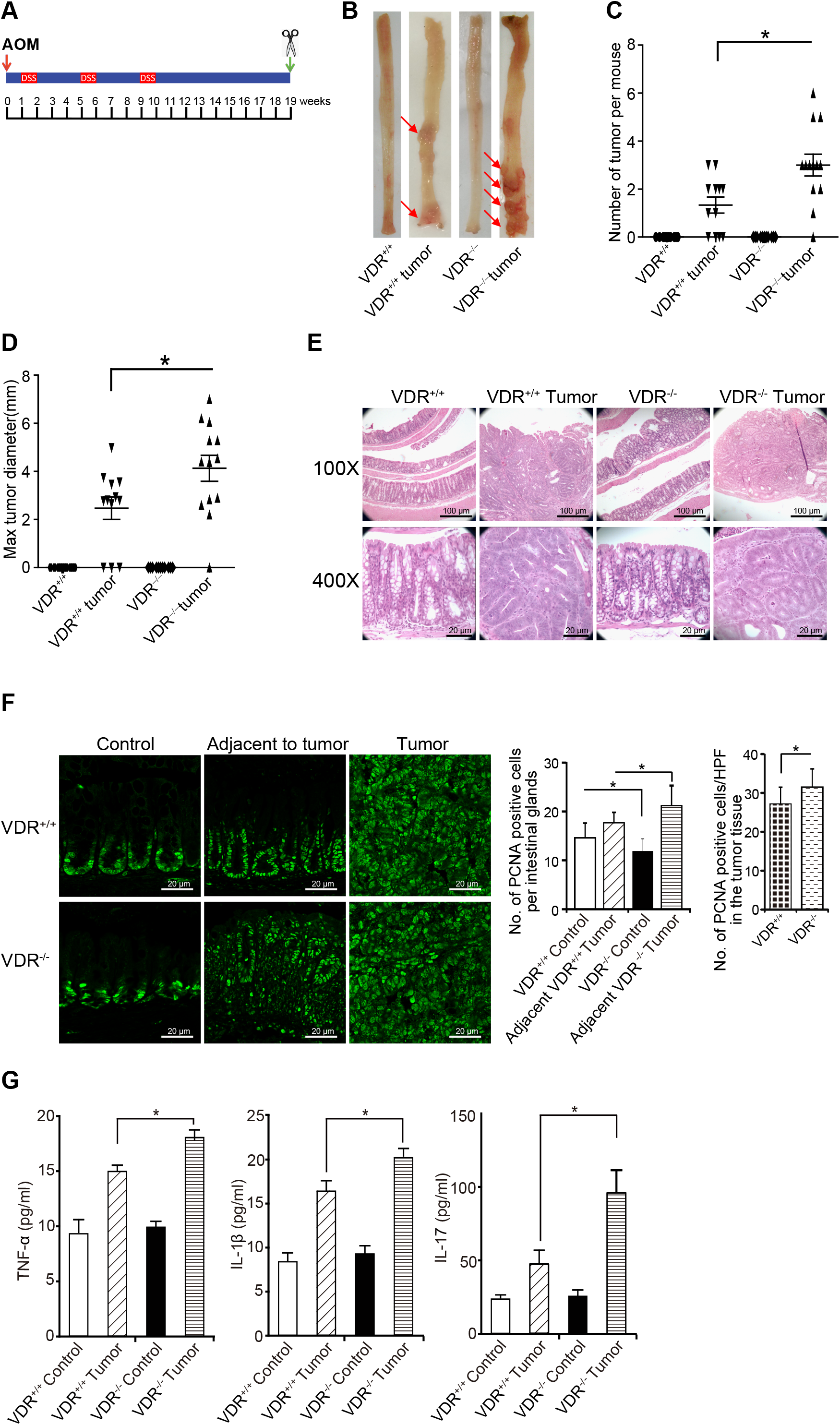
VDR^-/-^ mice developed a greater number of tumors compared to tumors in VDR^+/+^ mice. **(A)** Schematic overview of the AOM/DSS-induced colon cancer model. AOM (10 mg/kg) was injected on day 0. At Day 7, 2% DSS solution was administered to mice in drinking water. Seven days of DSS was followed by three weeks of drinking water that was free of DSS. An additional two cycles of DSS were administered prior to scarification at Week 19. **(B)** Colonic tumors *in situ*. Representative colons from different groups. Tumors are indicated by red arrows. **(C)** Tumor numbers in AOM-DSS-induced colon cancer model: VDR^+/+^ and VDR^-/-^ mice (data are expressed as mean ± SD. n = 10-13, one-way ANOVA test, *P < 0.05). **(D)** Max tumor size in AOM-DSS induced colon cancer model: VDR^+/+^ and VDR^-/-^ mice (data are expressed as mean ± SD. n = 10-13, one-way ANOVA test, *P < 0.05). **(E)** Representative H&E staining of “Swiss rolls” of representative colons from the indicated groups. Images are from a single experiment and are representative of 10 mice per group. **(F)** Quantitation of PCNA-positive cells in control mucosa per intestinal glands or in the tumors tissue per high-power field. PCNA expression in the tumor tissue of VDR^-/-^ mice was significantly higher, compared to that in the VDR^+/+^ mice (data are expressed as mean ± SD. n = 5, student’s t-test, *P < 0.05). **(G)** Serum cytokines such as TNF-α, IL-1β, and IL-17 were significantly increased, particularly in the AOM-DSS-induced VDR^-/-^ mice colon cancer model. Each single experiment was assayed in triplicate. Data are expressed as mean ± SD. n = 6, one-way ANOVA test, *P < 0.05.

### VDR deletion leads to decreased Claudin-5 expression in tumor tissues

We examined changes in barrier function by testing intestinal permeability in mice with or without tumors. Mice were gavaged with fluorescein dextran (molecular weight 3 kDa). After 4-h, blood samples were collected for fluorescence intensity measurements. Higher fluorescence intensity is indicative of higher intestinal permeability. As shown in **Fig. 3A**, AOM/DSS treatment induced increased intestinal permeability in both VDR^+/+^ and VDR^-/-^ mice, while the VDR^-/-^ mice exhibited significantly higher permeability post-treatment. Based on the *in vivo* intestinal permeability data, we hypothesized that the TJ proteins would be altered in the AOM/DSS mice. In the VDR^-/-^ mice, we observed significant downregulation of Claudin-5 at the mRNA and protein levels in the colon (**Fig. 3B & 3C**). Claudin-5 staining was observed at the crypt surface and at the lower portion of the intestine. Reduced Claudin-5 expression was confirmed through the immunostaining of AOM/DSS mice (**Fig. 3D & 3G**). However, VDR deletion did not alter the expression of the TJ protein Claudin-7 in the colon of VDR^-/-^ mice compared to that in VDR^+/+^ mice (**Fig. 3E**). VDR expression was also decreased in mice with AOM/DSS-induced colon cancer (**Fig. 3F and 3H**). Moreover, we used our recently established method to measure VDR levels according to qPCR in fecal samples [29]. We detected a significant reduction in VDR in fecal samples from mice with tumors (**Fig. 3I**). These data also suggest a decreased VDR in epithelial cells that are shed from mice with tumors.

**Fig. 3.**
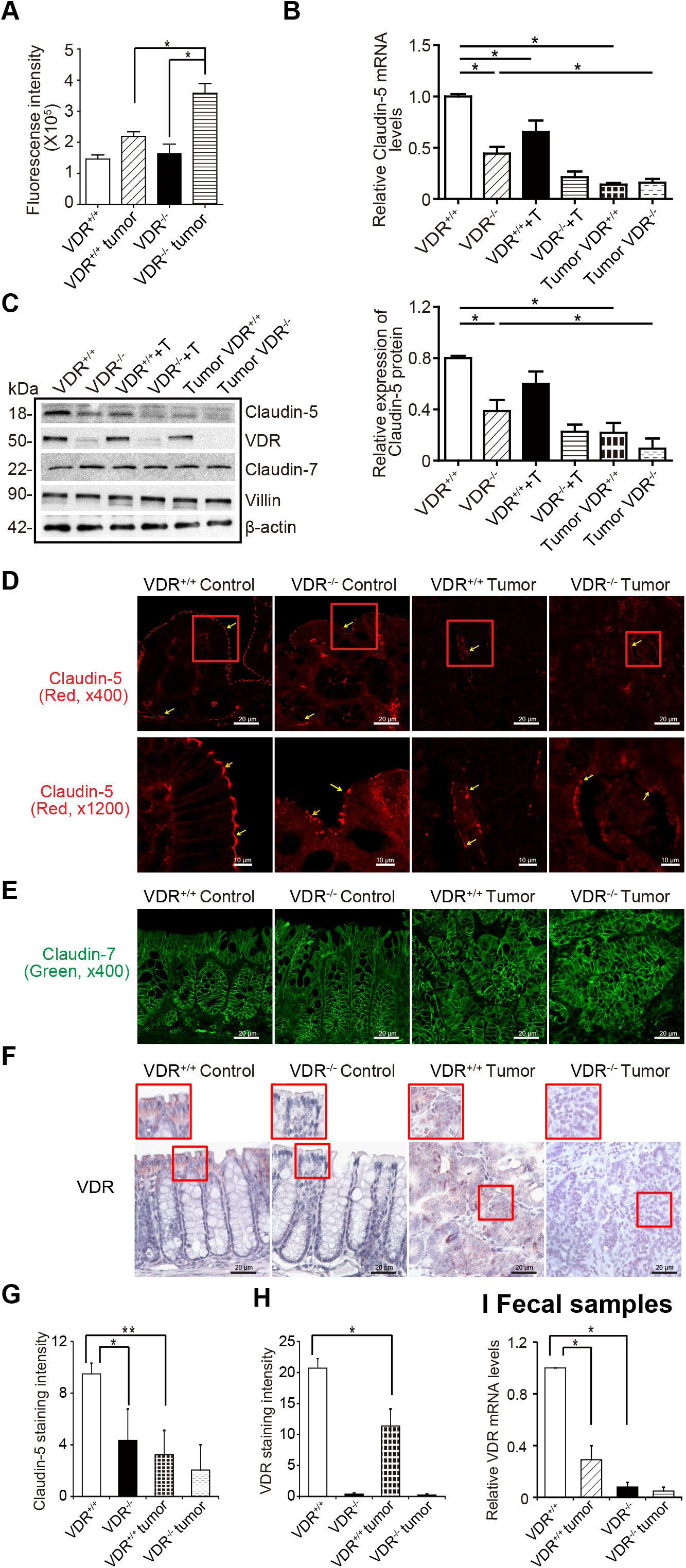
VDR deletion led to decreased Claudin-5 expression in tumor tissues. **(A)** Intestine permeability increased in the AOM-DSS-induced VDR^-/-^ mice colon cancer model. Fluorescein Dextran (Molecular weight 3 kDa, diluted in HBSS) was gavaged (50 mg/g mouse). Four hours later, mouse blood samples were collected for fluorescence intensity measurement (data are expressed as mean ± SD; n = 5 mice/group, 1-way ANOVA test; \**P* < 0.05). **(B)** VDR deletion decreased Claudin-5 at the mRNA level in the colon (data are expressed as mean ± SD. n = 5, one-way ANOVA test, *P < 0.05). **(C)** VDR deletion decreased Claudin-5 at the protein level in the colon (data are expressed as mean ± SD. n = 5, one-way ANOVA test, *P < 0.05). (**D**) **(G)** Claudin-5 was decreased in the tumor tissue of VDR^-/-^ mice, compared to levels in the tumor tissue of VDR^+/+^ mice according to immunofluorescence staining. Images are from a single experiment and are representative of 6 mice per group. (Data are expressed as mean ± SD. n = 6, one-way ANOVA test, *P < 0.05). **(E)** Claudin-7 was unchanged in the tumor tissue of VDR^-/-^ mice compared to levels in the tumor tissue of VDR^+/+^ mice according to immunofluorescence staining. Images are from a single experiment and are representative of 6 mice per group. **(F)(H)** Intestinal VDR expression was decreased in the AOM-DSS-induced colon cancer model. Images are from a single experiment and are representative of 6 mice per group. Data are expressed as mean ± SD. n = 6, one-way ANOVA test, *P < 0.05). **(I)** VDR levels in fecal samples were detected using RT-PCR. VDR expression was downregulated in the AOM-DSS-treated VDR^+/+^ mice (data are expressed as mean ± SD. *n* = 5, one-way ANOVA test, *P < 0.05).

### Conditional deletion of intestinal epithelial VDR led to increased permeability and reduced Claudin-5 in the AOM/DSS cancer model

Intestine permeability was also significantly increased in VDR^ΔIEC^ mice with conditional deletion of intestinal epithelial VDR (**Fig. 4A**). Intestinal epithelial VDR-specific deletion led to significantly decreased Claudin-5 at the mRNA level in the colon (**Fig. 4B**) and further decreased in the mice with colon cancer; however, other Claudin, such as Claudin-7, was not altered in the absence of VDR. At the protein level, we found the reduced Claudin-5 in the VDR^ΔIEC^ mice (**Fig. 4C**). In tumor tissues of VDR^ΔIEC^ mice epithelial Claudin-5 was disorganized (**Fig. 4D)** and significantly decreased, compared to that in tumors of VDR^loxp^ mice (**Fig. 4F**). In contrast, Claudin-7 was not altered in tumors from VDR^ΔIEC^ mice compared to the tumor tissue of VDR^loxp^ mice (**Fig. 4E**). The VDR expression in fecal samples was downregulated in the AOM-DSS VDR^loxp^ mice (**Fig. 4G**).

**Fig. 4.**
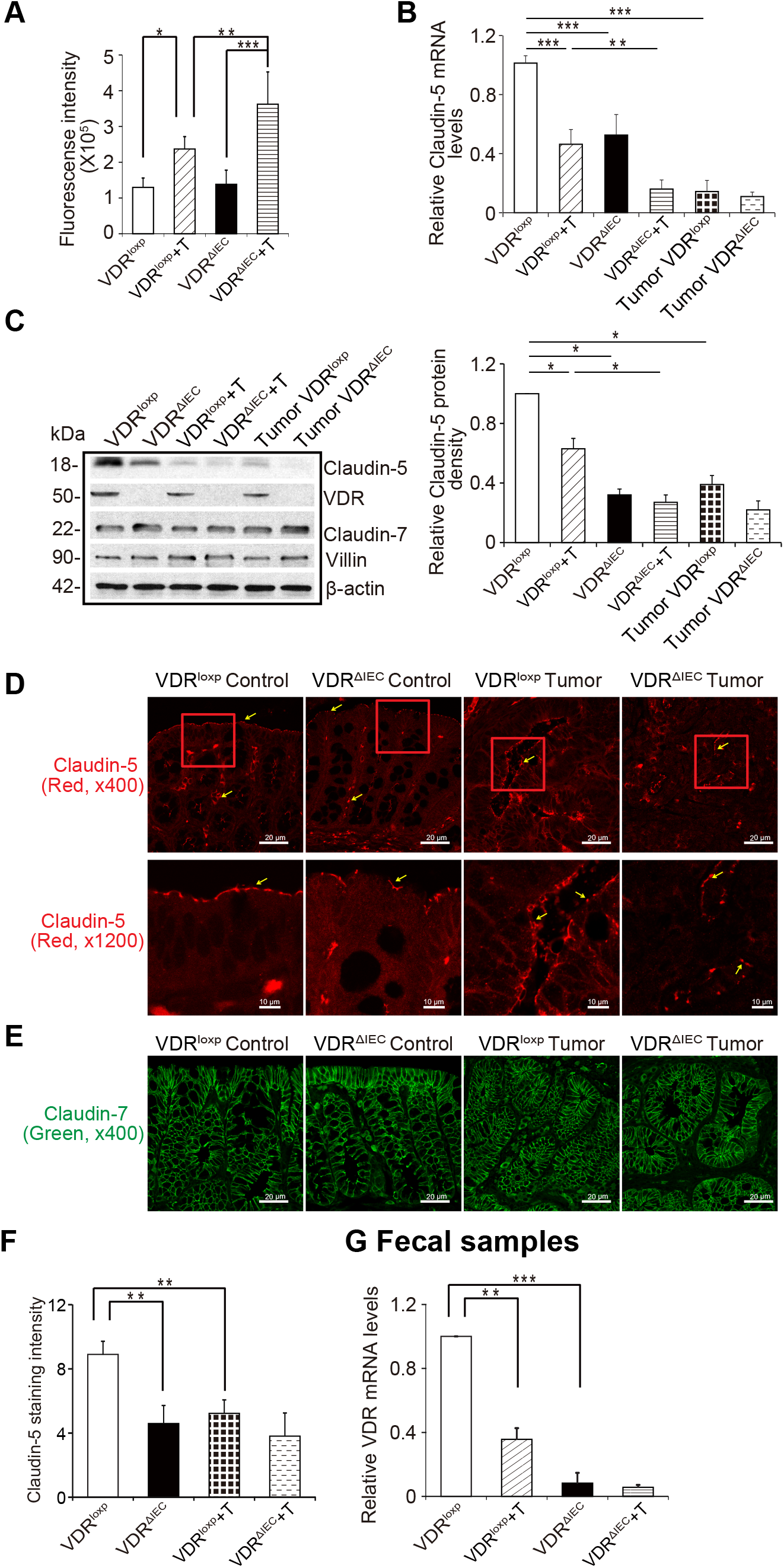
VDR-specific deletion in mouse intestines lead to decreased Claudin-5 expression in tumor tissues. **(A)** Intestine permeability was increased in the AOM-DSS-induced VDR^ΔIEC^ mice colon cancer model (data are expressed as mean ± SD; n = 5 mice/group, 1-way ANOVA test; \**P* < 0.05). **(B)** VDR-specific deletion in mouse intestines decreased Claudin-5 at the mRNA level in the colon (data are expressed as mean ± SD. n = 5, one-way ANOVA test, *P < 0.05). **(C)** VDR-specific deletion in mouse intestines decreased Claudin-5 protein in the colon (data are expressed as mean ± SD. n = 5, one-way ANOVA test, *P < 0.05). **(D)** Claudin-5 was decreased in the tumor tissue of VDR^ΔIEC^ mice compared to levels in the tumor tissue of VDR^loxp^ mice according to immunofluorescence staining. Images are from a single experiment and are representative of 6 mice per group. **(E)** Claudin-7 expression was not changed in the AOM-DSS-induced VDR^loxp^ mice colon cancer model. Images are from a single experiment and are representative of 6 mice per group. (**F**) Intensity of the staining of Claudin-5. (Data are expressed as mean ± SD. n = 6, one-way ANOVA test, *P < 0.05). **(G)** VDR level in fecal samples was detected by RT-PCR. VDR expression was downregulated in the AOM-DSS-treated VDR^loxp^ mice (data are expressed as mean ± SD. *n* = 3, one-way ANOVA test, *P < 0.05).

### Identification of the Vitamin D-response element (VDRE) in the Claudin-5 promoter

To confirm the direct regulation of VDR on Claudin-5, we examined various models at the basal level without any treatment *in vivo* and *in vitro*. In the VDR^-/-^ mice, we observed that these mice possessed lower Claudin-5 protein levels in the colon than did VDR^+/+^ mice, and TJ Claudin-7 was not altered in the absence of VDR (**Fig. S1A**). We further detected significantly decreased mRNA levels of Claudin-5 in the intestines of VDR^-/-^ mice (**Fig. S1B**). The density of Claudin-5 fluorescence staining was weaker in the VDR^-/- mouse^ intestines (**Fig. S1C**). We also assessed the specificity of intestinal VDR in regulating Claudin-5 expression in VDR^ΔIEC^ mice (**Fig. S1D**). Claudin-5 mRNA levels were significantly reduced in VDR^ΔIEC^ mice compared to the levels in VDR-lox mice (**Fig. S1E**). As expected, Claudin-7 expression remained unchanged. These data indicate that intestinal VDR specifically regulates the expression level of Claudin-5 in the colon. To confirm our findings *in vitro*, we used MEFs with VDR deletion. Lack of VDR led to a robust decrease in Claudin-5 protein and mRNA levels in VDR^-/-^ MEFs at the basal level (**Fig. S1F** and **S1G**). The density of Claudin-5 fluorescence staining was also weaker in VDR^-/-^ MEFs (**Fig. S1H**).

VDR acts as a transcription factor to regulate the expression of its target genes [30, 31]. Activated VDR binds to VDRE in the target gene promoter to regulate gene transcription [32]. We reasoned that VDR may bind to the *Claudin-5 promoter* to thus alter the mRNA expression of the Claudin-5 gene. Further, we performed a ChIP assay using the colon mucosal extract from VDR^-/-^ mice and nonspecific IgG as a negative control to assess the binding of VDR to the Claudin-5 promoter. The samples were amplified by conventional PCR using Ikβα as a positive control and Claudin-1 as a negative control as indicated in previous publications [33]. CHIP-PCR demonstrated that VDR bound to the Claudin-5 promoter in the VDR^+/+^ mouse colon **(Fig. 5A)**. The VDRE sequence (AGTTCAAGTGGTTCT) within the Claudin-5 promoter region is shown in Figure **5B**. However, siRNA-based Claudin-5 knock-down did not reduce VDR expression at the mRNA level (**Fig. 5C)**. At the protein level, reduced Claudin-5 did not change the status of VDR protein or Claudin-7 at the protein level (**Fig. 5D)**. These results suggest that VDR transcriptionally regulates Claudin-5 at the mRNA level and that VDR is the upstream regulator of Claudin-5.

**Fig. 5.**
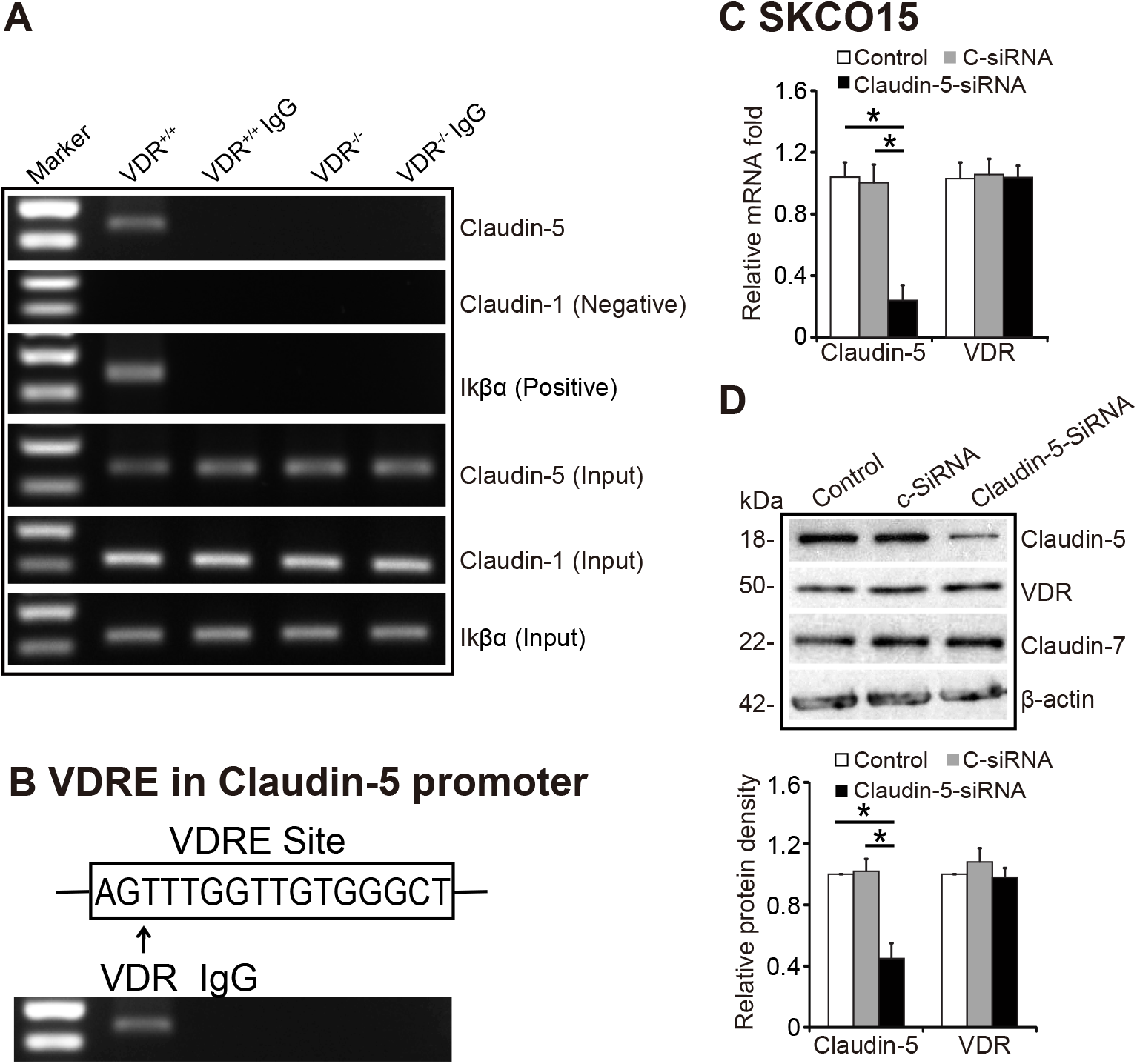
VDR binds to the Claudin-5 promoter *in vivo* and *in vitro*. **(A)** CHIP-PCR amplification demonstrated that VDR binds to the promoter regions of Claudin-5 in mouse colons. PCR assays were performed and included input-positive controls and IgG/villin-negative controls. n = 3 separate experiments. **(B)** Claudin-5 promoter regions with VDRE sequence. (**C**) Claudin-5 knockdown using siRNA (40 nM for 72 hours) did not reduce VDR expression at the mRNA level. (Data are expressed as mean ± SD. *n* = 3, one-way ANOVA test, *P < 0.05). (**D**) The protein expression in SKCO15 cells using siRNA (40 nM for 72 hours). (data are expressed as mean ± SD. *n* = 3, one-way ANOVA test, *P < 0.05).

### High VDR levels led to increased Claudin-5 protein and mRNA levels *in vitro*

We then explored the possibility of enhancing VDR to maintain the physiological level of Claudin-5. Vitamin D3 is known to increase VDR expression and to activate VDR signaling. We used the human colonic epithelial SKCO15 cell line that is widely used to study TJs [34, 35]. Claudin-5 mRNA level was significantly elevated in SKCO15 cells treated with vitamin D3, while Claudin-7 mRNA was not altered by vitamin D3 treatment (**Fig. 6A)**. The protein level of Claudin-5 was induced by vitamin D3 (**Fig. 6B**). *In vivo*, Claudin-5 mRNA levels were also increased in vitamin D3-treated mice (**Fig. 6C**). Colonoids are three-dimensional (3D) cell cultures that incorporate a number of key features of the colon [36]. In this study, we developed human colonoids (**Fig. 6E**), and we observed vitamin D3 treatment significantly increased Claudin-5 mRNA levels in these colonoids (**Fig. 6F**). Furthermore, vitamin D3 treatment significantly increased Claudin-5 protein levels in human colonoids, whereas there was no change of Claudin-7 after vitamin D3 treatment (**Fig. 6G**).

**Fig. 6.**
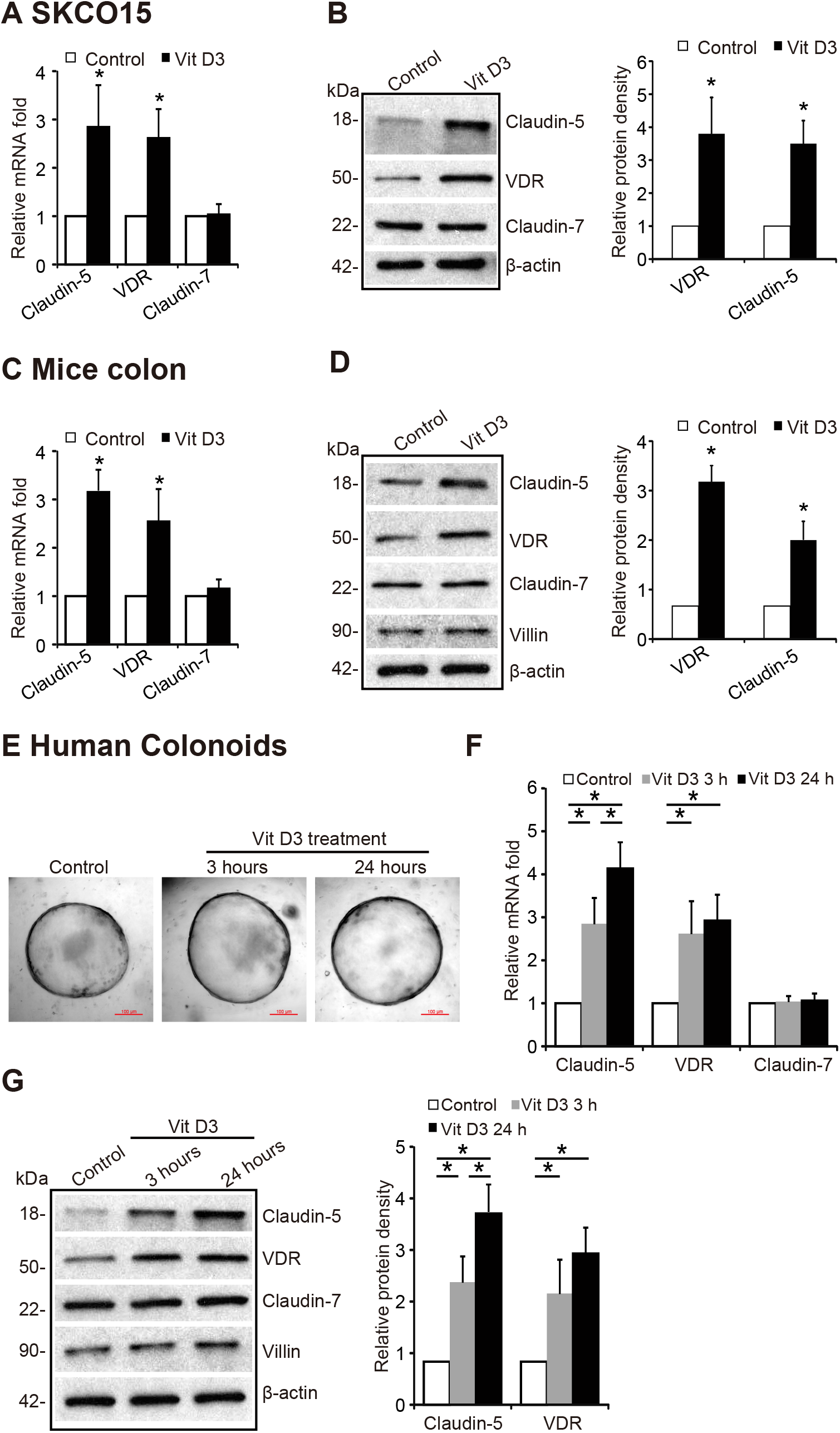
High VDR levels led to increased Claudin-5 at the protein and mRNA levels *in vitro*. **(A)** Claudin-5 mRNA and (**B**) protein levels were increased after 24-hour vitamin D3 treatment at 20 nM in SKCO15 cells (data are expressed as mean ± SD. student’s t-test, *P < 0.05. n = 3). **(C)** Claudin-5 mRNA and **(D)** protein levels were higher in vitamin D3-treated VDR^+/+^ mice. VDR^+/+^ mice (6-8 weeks) were gavaged by 0.2 μg vitamin D3 in 0.1 ml corn oil for 3 times per week for 4 weeks (data are expressed as mean ± SD. student’s t-test, *P < 0.05. n= 5 mice / group). **(E)** The micrographs show representative human colonoids that were treated with Vit D_3_ (20 nM) for the indicated time points. **(F)** Claudin-5 mRNA and (**G**) protein levels were increased after vitamin D3 treatment in human colonoids (data are expressed as mean ± SD, n= 5, one-way ANOVA test, *P < 0.05).

### Intestinal epithelial VDR overexpression protected against the loss of Claudin 5 in respond to inflammation

To further study the protective role of VDR in_maintaining TJs in inflammation, we generated a conditional intestinal epithelial VDR specific-overexpressed (O-VDR) mouse model. Epithelial VDR overexpression in mouse intestines significantly increased Claudin-5 expression at both the mRNA and protein levels (**Fig. 7A & B**). Claudin-5 exhibited a less decreaseat the mRNA and protein and mRNA levels in the colon of O-VDR mice treated with DSS, compared to that in the O-VDR^loxp^ mice (**Fig. 7C & 7D).** Using IF staining, we determined that Claudin-5 was better preserved in the colon of O-VDR mice treated with DSS compared to that in the O-VDR^loxp^ mice **(Fig. 7E & 7G).** As anticipated, Claudin-7 expression was unchanged in the intestinal tissue of O-VDR mice that were treated with DSS compared to that in the O-VDR^loxp^ mice **(Fig. 7F).** VDR levels in fecal samples were detected using RT-PCR. VDR level was less decreased in DSS-treated O-VDR mice, compared to that in O-VDR^loxp^ mice treated with DSS **(Fig. 7H)**. Moreover, there were fewer inflammatory cytokines such as IL-1β and IL-17 in the colons of DSS-induced O-VDR mice compared to that in O-VDR^loxp^ mice **(Fig. 7I)**.

**Fig. 7.**
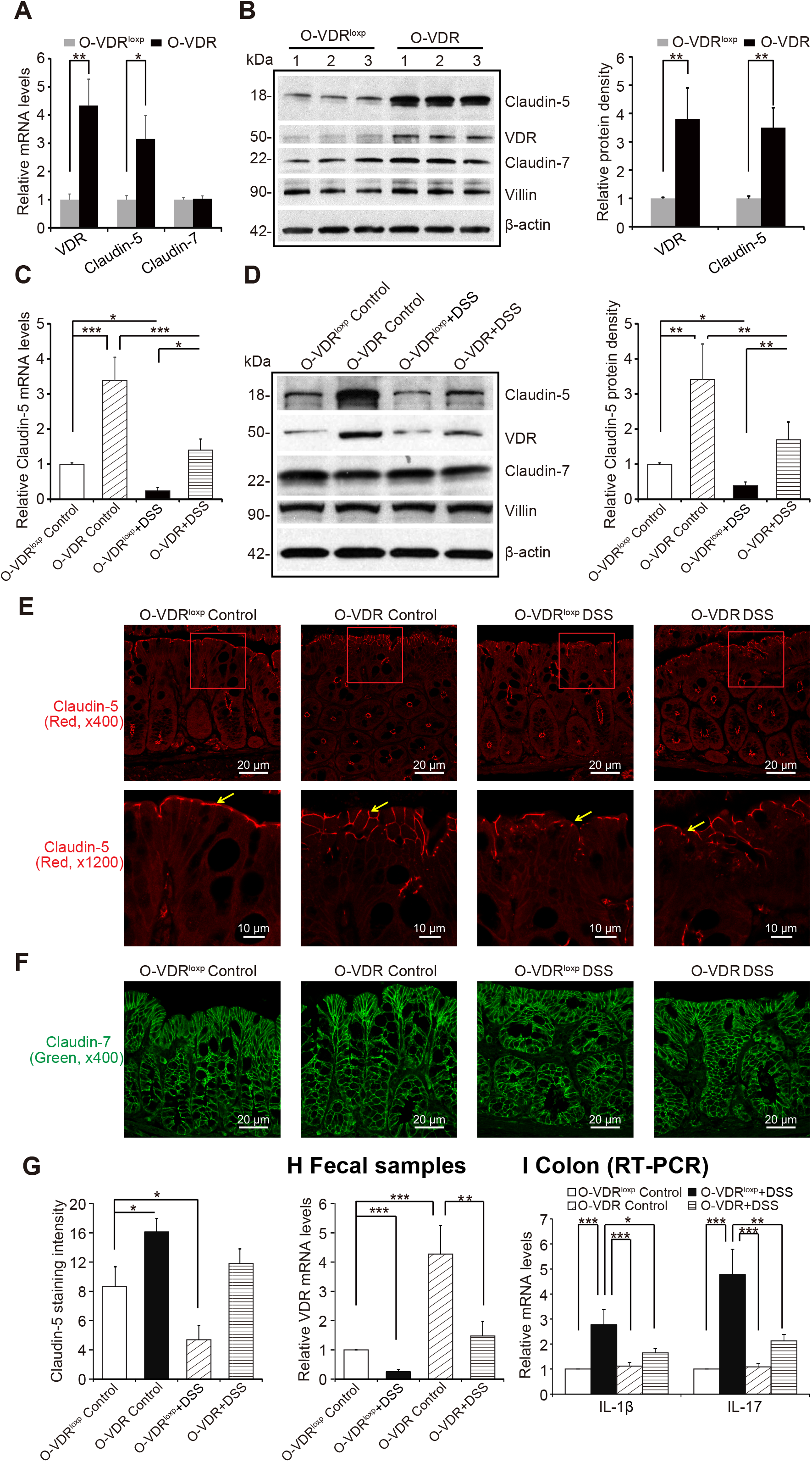
Overexpressed intestinal epithelial VDR led to increased Claudin-5 and reduced inflammation *in vivo*. **(A)** VDR overexpression in mice intestines increased Claudin-5 expression in the colon at the mRNA and (B) protein levels (data are expressed as mean ± SD. n = 3, one-way ANOVA test, *P < 0.05). **(C)** Claudin-5 was less decreased at the mRNA and (**D**) protein levels in the intestinal tissue of O-VDR mice treatment with DSS compared to levels in the O-VDR^loxp^ mice (data are expressed as mean ± SD. n = 3, one-way ANOVA test, *P < 0.05). **(E)** Claudin-5 was less decreased in the intestinal tissue of O-VDR mice treated with DSS compared to levels in the O-VDR^loxp^ mice, according to immunofluorescence staining. Images are from a single experiment and are representative of 5 mice per group. **(F)** Claudin-7 was unchanged in the intestinal tissue of O-VDR mice treated with DSS compared to levels in the O-VDR^loxp^ mice according to immunofluorescence staining. Images are from a single experiment and are representative of 5 mice per group. **(G)** Intensity of the staining of Claudin-5. Images are from a single experiment and are representative of 5 mice per group. (Data are expressed as mean ± SD. n = 5, one-way ANOVA test, *P < 0.05). **(H)** VDR level in fecal samples was detected by RT-PCR. VDR expression was less decreased in O-VDR mice treated with DSS compared to levels in the O-VDR^loxp^ mice treated with DSS (data are expressed as mean ± SD. *n* = 3, one-way ANOVA test, *P < 0.05). **(I)** The inflammatory cytokines IL-1β and IL-17 were less increased in the DSS-induced O-VDR mice colitis model compare to levels in O-VDR^loxp^ mice (data are expressed as mean ± SD. *n* = 3, one-way ANOVA test, *P < 0.05).

## Discussion

In the current study, we determined that low colonic VDR expression was significantly correlated with the reduction of Claudin-5 in human CRC, and we demonstrated that VDR is important for the maintenance of cellular and physiological levels of TJ protein Claudin-5 in the colon to prevent inflammation and tumorigenesis. Our study further revealed a complex role for vitamin D/VDR regulation of *CLDN-5* in the development of colon cancer. Lack of VDR led to a reduction of Claudin-5 in tumors, and enhancing VDR increased Claudin-5 to protect the intestinal epithelial cells from tumorigenesis. At the molecular level, our data have demonstrated that the *CLDN-5* gene is a newly discovered downstream target of the transcriptional factor VDR. Overall, we noted a link between VDR signaling and barrier functions in CRC, thus suggesting a potential biomarker and target for a novel therapeutic strategy. Our study provides insight into how VDR signaling is involved in the tissue barrier related to tumorigenesis.

The intestinal barrier includes several elements that aid in its function as a physical and immunological barrier. These elements include the intestinal microbiota, secretory immunoglobulin A, antimicrobial peptides, the inner lamina propria, and epithelial cells. At the cellular level, epithelial cells play physical and physiological roles in health and disease. VDR signaling is involved in the epithelial barrier function related to various human diseases and remains largely unexplored [37]. As a nuclear receptor, VDR mediates most known functions of 1,25-dihydroxyvitamin D (1,25(OH)_2_D_3_), the active form of vitamin D [4]. However, the role of VDR has rarely been evaluated in studies examining human colon cancer. A recent study among patients with digestive tract cancer and vitamin D supplementation determined that when compared to placebo, this treatment did not result in significant improvement in relapse-free survival at 5 years [38]. The dosage of vitamin D3 was insufficient among participants who possessed more severe deficiencies at baseline. Therefore, the status of the VDR level must to be considered over the course of many trials or as a biological measurement to clarify the underlying mechanisms. The traditional model of treatment using vitamin D that guided early vitamin studies should give way to a model incorporating more complex mechanisms of action of the vitamin D/VDR system. The intestinal barrier has been investigated by various methods, but correlation of results across studies is difficult, representing a major shortcoming in the field [39].

The current study provides important insights into how VDR regulates Claudin-5 expression under normal physiological conditions and during tumor growth in the colon. We revealed a positive correlation between VDR and Claudin-5 at the mRNA and protein levels in healthy and tumor colons, thus suggesting the unique role of Claudin-5 in the intestine. There are 27 claudin family members that contribute to tight junctions [40], and not all claudins are the same. Claudin-2- and Claudin-12 form paracellular Ca^2+^ channels in intestinal epithelia and are important for vitamin D-dependent calcium homeostasis [41]. Our previous studies have shown that Claudin-2 is hyperregulated in colitis with VDR reduction [42, 43]. Our current study has demonstrated the mechanism on VDR-dependent function of Claudin-5 in the intestine. Interestingly, we found that the tight junction Claudin-7 was not altered in response to VDR-deficient status in the colon. In the lungs, VDR may play an important role in maintaining the pulmonary barrier integrity. We have reported that VDR deletion could increase lung permeability by altering the expression of TJ molecules, particularly Claudin-2, −4, −10, −12, and −18 [44]. Abnormal gut barrier function may serve as a biomarker for the risk of IBD onset [45]. Our findings also suggest that the positively correlated status of VDR and Claudin-5 could be potentially applied to risk assessment, early detection, and prevention of CRC, including colitis-associated colon cancer.

Colorectal cancer is the second-leading cause of cancer-related death and is most curable in its early stages. Remarkable progress has been made in regard to colon cancer therapy, including targeting barrier functions and microbiome [46]. The anti-TNF era has revealed that mucosal healing is a key goal for therapy that predicts clinical remission and resection-free survival in cancer patients. Many new targets (e.g., Jak inhibition, Toll-like receptor 9 stimulation, and the addressin mucosal vascular addressin cell adhesion molecule 1 emerge) have been recently tested for induction of mucosal healing and protection and for induction and maintenance of remission in IBD. We aimed to provide a detailed understanding of how VDR status contributes to changes in TJs in the context of intestinal inflammation and colon cancer. Currently, there are no guidelines for monitoring vitamin D status, treating hypovitaminosis D, and maintaining optimal vitamin D stores in patients with IBD [47] or in CRC. These tasks may prove particularly difficult due to malabsorption, gastrointestinal losses, and increased permeability that are associated with intestinal dysfunction. Based on the research progress regarding the novel roles of VDR in intestinal immunity and barrier functions, we expect that studies on VDR in intestinal barriers of colitis and colon cancer will have a marked impact on the prevention, diagnosis, and therapy of colitis and colon cancer patients.

## Materials and Methods

### Human tissue samples

This study was performed in accordance with approval from the University of Rochester Ethics Committee (RSRB00037178) and UIC Ethics Committee (Institutional Review Board: 2017-0384). Colorectal tissue samples were obtained from 10 CRC patients with neoplasia and 10 patients without neoplasia patients (49–74years old). Human tissues for organoids are from healthy volunteers.

### Gene expression datasets

For expression analyses, we used microarray data reported in the NCBI Gene Expression Omnibus database (GEO). To find the correlation between VDR and Claudin-5 at the gene expression level, we gathered data by searching the Gene Expression Omnibus (https://www.ncbi.nlm.nih.gov/geo/) for expression profiling studies using colonic samples from colon cancer subjects. We randomly identified the GEO database reference series: GSE4183 [48], GSE8671 [49], GSE10714 [50] and GSE37283) [51]. In these studies, the authors performed microarray analysis using colonic biopsy samples from healthy controls as well as from the inflamed and non-inflamed colonic mucosa from CRC subjects. From the databases, 40 healthy controls and 62 CRC patients were randomly selected for CRC group, while 16 healthy controls and 18 Colitis-associated CRC were randomly selected for colitis-associated group. Both were subjected to further analyses.

### Animals

VDR^-/-^ mice on a C57BL/6 background were obtained by breeding heterozygous VDR^+/-^mice[43]. VDR^ΔIEC^ mice were obtained by crossing the VDR^LoxP-B^ mice, originally provided by Dr. Geert Carmeliet, with villin-cre mice (Jackson Laboratory, 004586), as we previously reported [52].

Intestinal-specific VDR-overexpressing (O-VDR) mice were generated in C57BL/6 mice strain background. The mouse VDR (mVDR) sequence was cloned into the Stbl3 vector (size 6631 bp). The mVDR was cloned in (from ~2210 bp to ~3316bp) under EF1A promoter (1bp to 1105bp). A LoxP site was integrated after EF1A promoter (from 1105 bp to 2210 bp). VDR expression in O-VDR mice is Cre driven [53]. This O-VDR^loxp^ mouse line is labeled as O-VDR^loxp^ in our gain of function study to distinct from the VDR^loxP/loxP^ mouse made for VDR^ΔIEC^ mice.

Experiments were performed on 2–3 months old mice including male and female. Mice were provided with water ad libitum and maintained in a 12 h dark/light cycle. The animal work was approved by the Rush University Animal Resources committee and UIC Office of Animal Care. The animal work was approved by the UIC Office of Animal Care (ACC 15-231,17-218, and 18-216).

### Induction of colon cancer by AOM-DSS in mice

Mice were treated with 10mg/kg of AOM (Sigma-Aldrich, Milwaukee, WI, USA) by intraperitoneal injection as previously described [24]. After a 7-day recovery period, mice received three cycles of 2% DSS in the drinking water. Tumor counts and measurements were performed in a blinded fashion under a stereo-dissecting microscope (Nikon SMZ1000, Melville, NY, USA). Microscopic analysis was performed for severity of inflammation and dysplasia on hematoxylin and eosin-stained ‘Swiss rolled’ colons by a gastrointestinal pathologist blinded to treatment conditions. Mice were scarified under anaesthesia.

### Induction of colitis and experimental design

Eight-to ten-week-old mice of a specific genetic background were grouped randomly into control and DSS treatment groups. Colitis was induced by adding 5% (weight/volume) dextran sodium sulfate (DSS) (mol. wt 36-50 kD; USB Corporation, Cleveland, OH, USA) to the drinking water for 7 days. Mice were monitored regularly, and their body weights were noted every day. All mice were provided a regular chow diet ad libitum. We checked the effect of DSS on both OVDR mice and compared them with their respective control group. On day 7, mice were sacrificed, and intestinal tissue and blood samples were harvested for RNA, protein, immunofluorescence, and cytokine analysis as described in the results section. The samples were immediately frozen and kept at −80°C until use.

### Vitamin D_3_ treatment *in vivo*

C57/BL/6 wild-type mice (6-8-week-old males and females) were gavaged with 1,25 D_3_ (0.2 μg/day in 100 μl of corn oil) 3 times a week for 4 weeks, as described in our previous study [54]. Intestinal tissue was collected following euthanasia.

### Cell culture

Mouse embryonic fibroblasts (MEF) were isolated from embryonic day 13.5 embryos generated from VDR^+/-^ x VDR^+/-^ mouse breeding as previously described (32). VDR^+/+^ and VDR^-/-^ MEFs were used in experiments after more than 15 passages when they had been immortalized. MEFs and SKCO15 cells were grown in high glucose Dulbecco’s Modified Eagle Medium (DMEM) (Hyclone, SH30243.01) containing 10% (v/v) fetal bovine serum (GEMINI, 900-108), 50 μg/ml streptomycin, and 50 U/ml penicillin (Mediatech, Inc., 30-002CI), as previously described [55].

### Colonoids cultures and treatment with Vit D3

Human colonoids were prepared and maintained as previously described [56]. Mini gut medium (advanced DMEM/F12 supplemented with HEPES, L-glutamine, N2, and B27) was added to the culture, along with R-Spondin, Noggin, EGF, and Wnt-3a. At day 7 after passage, colonoids were treated by Vit D3 (20 nM) for indicated times.

### Western blot analysis and antibodies

Mouse colonic epithelial cells were collected by scraping the tissue from the colon of the mouse, including the proximal and distal regions [52]. The cells were sonicated in lysis buffer (10 mM Tris, pH 7.4, 150 mM NaCl, 1 mM EDTA, 1 mM EGTA, pH 8.0, 1% Triton X-100) with 0.2 mM sodium ortho-vanadate, and protease inhibitor cocktail. The protein concentration was measured using the BioRad Reagent (BioRad, Hercules, CA, USA). Cultured cells were rinsed twice with ice-cold HBSS, lysed in protein loading buffer (50 mM Tris, pH 6.8, 100 mM dithiothreitol, 2% SDS, 0.1% bromophenol blue, 10% glycerol), and then sonicated. Equal amounts of protein were separated by SDS-polyacrylamide gel electrophoresis, transferred to nitrocellulose, and immunoblotted with primary antibodies. The following antibodies were used: anti-Claudin-5 (Invitrogen, 35-2500, Carlsbad, CA, USA), anti-Claudin-7 (Invitrogen, 34-9100, Carlsbad, CA, USA), anti-VDR (Santa Cruz Biotechnology, SC-13133, Dallas, TX, USA), anti-Villin (Santa Cruz Biotechnology, SC-7672 Dallas, TX, USA), or anti-β-actin (Sigma-Aldrich, A5316, St. Louis, MO, USA) antibodies and were visualized by ECL (Thermo Fisher Scientific, 32106, Waltham, MA, USA). Membranes that were probed with more than one antibody were stripped before re-probing.

### Immunofluorescence

Colonic tissues were freshly isolated and embedded in paraffin wax after fixation with 10% neutral buffered formalin. Immunofluorescence was performed on paraffin-embedded sections (4 μm), after preparation of the slides as described previously [52], [55] followed by incubation for 1 hour in blocking solution (2% bovine serum albumin, 1% goat serum in HBSS) to reduce nonspecific background. The tissue samples were incubated overnight with primary antibodies at 4°C. The following antibodies were used: anti-Claudin-5 and anti-Claudin-7. Slides were washed 3 times for 5 minutes each at room temperature in wash buffer. Samples were then incubated with secondary antibodies (goat anti-rabbit Alexa Fluor 488, Molecular Probes, CA; 1:200) for 1 hour at room temperature. Tissues were mounted with SlowFade Antifade Kit (Life technologies, s2828, Grand Island, NY, USA), followed by a coverslip, and the edges were sealed to prevent drying. Specimens were examined with a Zeiss laser scanning microscope LSM 710 (Carl Zeiss Inc., Oberkochen, Germany).

### Immunohistochemistry (IHC)

After preparation of the slides, antigen retrieval was achieved by incubation of the slides for 15 min in the hot preheating sodium citrate (pH 6.0) buffer, and 30 min of cooling at room temperature. Endogenous peroxidases were quenched by incubating the slides in 3% hydrogen peroxide for 10 min, followed by three rinses with HBSS, and incubation for 1 hour in 3% BSA + 1% goat serum in HBSS to reduce nonspecific background. Primary antibodies VDR (1:300) was applied for overnight in a cold room. After three rinses the slides with HBSS, they were incubated in secondary antibody (1:100, Jackson ImmunoResearch Laboratories, Cat.No.115-065-174, West Grove, PA, USA) for 1 hour at room temperature. After washing with HBSS for 10 minutes, the slides were incubated with vectastain ABC reagent (Vector Laboratories, Cat.No. PK-6100, Burlingame, CA 94010, USA) for 1 hour. After washing with HBSS for five minutes, color development was achieved by applying peroxidase substrate kit (Vector Laboratories, Cat.No. SK-4800, Burlingame, CA 94010) for 2 to 5 minutes, depending on the primary antibody. The duration of peroxidase substrate incubation was determined through pilot experiments and was then held constant for all of the slides. After washing in distilled water, the sections were counterstained with haematoxylin (Leica, Cat.No.3801570, Wetzlar, Germany), dehydrated through ethanol and xylene, and cover-slipped using a permount (Fisher Scientific, Cat.No.SP15-100, Waltham, MA, USA).

### Real Time quantitative PCR

Total RNA was extracted from epithelial cell monolayers or mouse colonic epithelial cells using TRIzol reagent (Fisher Scientific, 15596026, Waltham, MA, USA) [52]. RNA reverse transcription was done using the iScript cDNA synthesis kit (Bio-Rad Laboratories, 1708891) according to the manufacturer’s directions. The RT-cDNA reaction products were subjected to quantitative real-time PCR using the CFX96 Real time PCR detection system (Bio-Rad Laboratories, Hercules, CA, USA) and iTaq™ Universal SYBR green supermix (Bio-Rad Laboratories, 1725121, Hercules, CA, USA) according to the manufacturer’s directions. All expression levels were normalized to β-actin levels of the same sample. Percent expression was calculated as the ratio of the normalized value of each sample to that of the corresponding untreated control cells. All real-time PCR reactions were performed in triplicate. Primer sequences were designed using Primer-BLAST or were obtained from Primer Bank primer pairs listed in **Table 1**.

**Table 1:**
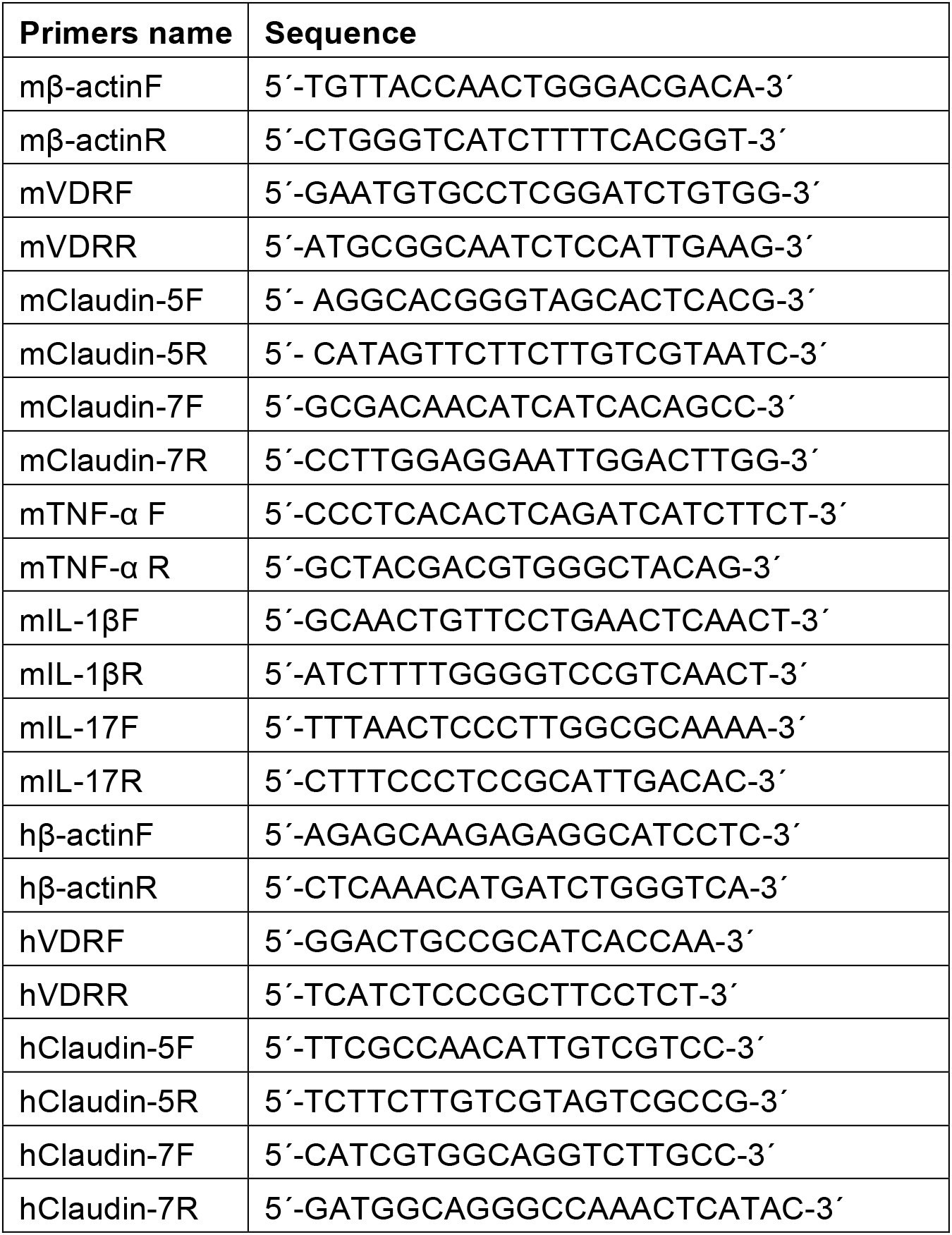
Real-time PCR Primers.

### Chromatin immunoprecipitation (CHIP) assay

Binding of VDR to the Claudin-5 promoter was investigated using the ChIP assay as described previously [57]. Briefly, mouse colonic epithelial cells were collected by scraping the tissue from the colon of the mouse, cells were treated with 1% formaldehyde for 10 min at 37°C. Cells were washed twice in ice-cold phosphate buffered saline containing protease inhibitor cocktail tablets. Cells were scraped into conical tubes, pelleted and lysed in SDS Lysis Buffer. The lysate was sonicated to shear DNA into fragments of 200–1000 bp (4 cycles of 10 s sonication, 10 s pausing, Branson Sonifier 250, USA). The chromatin samples were pre-cleared with salmon sperm DNA–bovine serum albumin-sepharose beads, then incubated overnight at 4 °C with VDR antibody. Immune complexes were precipitated with salmon sperm DNA-bovine serum albumin-sepharose beads. DNA was prepared by treatment with proteinase K, extraction with phenol and chloroform, and ethanol precipitation.

### Multiplex ELISA assay

A mouse-specific ProcartalPlex™ Multiplex Immunoassay (26) plate from Invitrogen Thermo Fisher Scientific was used to detect serum cytokine levels. The assay was performed using the manufacturer’s instruction manual using proper standards. Eventually, the plate was read using a Megpix Luminex machine.

### Test fecal VDR by PCR

Total RNA was extracted from mouse fecal samples, as previously described.[58] Briefly, About 100 mg of Frozen fecal pellet was used for RNA extraction by using Trizol Reagent (Thermo Fisher Scientific, Cat.No.15596018, Waltham, MA, USA). To increase RNA yield in high quality, RNeasy minispin column (Qiagen, Cat No.217004, Hilden, Germany) was used by following the manufacturer’s instructions.

### Statistical Analysis

All data are expressed as the mean ± SD. All statistical tests were 2-sided. All *p* values < 0.05 were considered statistically significant. After the Shapiro-Wilk test confirmed that the data are normally distributed, the differences between samples were analyzed using unpaired *t* test for two groups and using one-way ANOVA for more than two groups as appropriate, respectively. The *p* values in ANOVA analysis and generalized linear mixed models were adjusted using the Tukey method to ensure accurate results. Pairwise correlation analyses and scatter plots were conducted staining intensity changes between VDR protein and Claudin-5, using SAS version 9.4 (SAS Institute, Inc., Cary, NC, USA). Other statistical analyses were performed using GraphPad Prism 6 (GraphPad, Inc., San Diego, CA., USA).

## Author Contributions

YZ: acquisition, analysis, and interpretation of data, drafting of the manuscript, and statistical analysis, SG: assistance with western blots and TJ data. YX: Statistical analysis, microbiome data analysis, and manuscript drafting. REC: Provided human biopsies and clinical perspectives of CRC. JS: study concept and design, analysis and interpretation of data, writing of the manuscript for important intellectual content, obtaining funding, and study supervision.

## Acknowledgments

We would like to thank Dr. David Zhou for assisting with the CRC human samples, Drs. Shaoping Wu and Rong Lu for assisting with the AOM/DSS model, and Jason S. Xia for proofreading. The contents do not represent the views of the United States Department of Veterans Affairs or the United States Government.

---

**Fig. S1.**
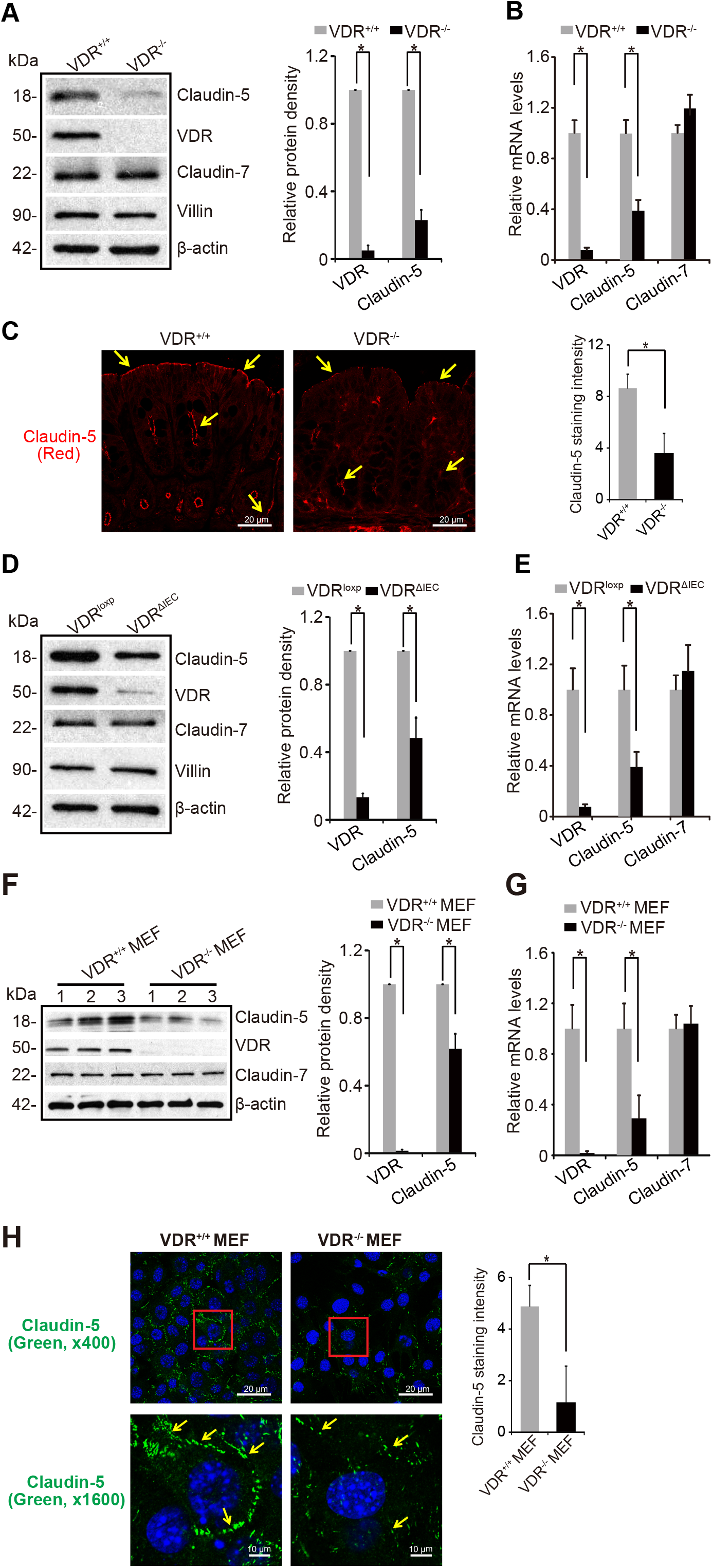
VDR deficiency in intestinal epithelial cells of mice leads to the reduction of Claudin-5 at both the mRNA and protein levels *in vivo*. **(A)** Claudin-5 protein and **(B)** mRNA levels were significantly lower in VDR^-/-^ mice compared to levels in the VDR^+/+^ mice (data are expressed as mean ± SD. n = 5, student’s t-test, *P < 0.05). **(C)** Location of Claudin-5 protein in the colons of VDR^+/+^ and VDR^-/-^ mice. Images are from a single experiment and are representative of 5 mice per group. **(D)** Claudin-5 protein and **(E)** mRNA levels were significantly lower in VDR^ΔIEC^ mice compared to levels in the VDR^loxp^ mice (data are expressed as mean ± SD. n = 5, student’s t-test, *P < 0.05). **(F)** Claudin-5 protein and **(G)** mRNA were both decreased in VDR^-/-^ MEF cells (data are expressed as mean ± SD. n = 3, student’s t-test, *P < 0.05). **(H)** Location and quantification of Claudin-5 protein in VDR^+/+^ and VDR^-/-^ MEF cells. Images are from a single experiment performed in triplicate. (Data are expressed as mean ± SD. n = 3, one-way ANOVA test, *P < 0.05).

